# Integrative CUT&Tag/RNA-Seq analysis of histone variant macroH2A1-dependent orchestration of human iPSCs reprogramming

**DOI:** 10.1101/2022.09.30.510248

**Authors:** Niccolò Liorni, Alessandro Napoli, Stefano Castellana, Sebastiano Giallongo, Daniela Řeháková, Oriana Lo Re, Irena Koutná, Tommaso Mazza, Manlio Vinciguerra

## Abstract

Human-induced pluripotent stem cells (iPSCs) can be derived from adult stem cells by forced expression of defined transcription factors. This paves the way for autologous iPSC-derived therapies, which, however, are not yet considered safe. Moreover, reprogramming of somatic cells into iPSCs is an inefficient process, in the range of 0.1%–1%. The epigenetic mechanisms implicated in iPSCs reprogramming are not well understood. The substitution of canonical histone H2A with macroH2A1 histone variant exon-spliced isoforms (macroH2A1.1 and macroH2A1.2) appears as an emerging regulator of iPSCs identity. In particular, we have previously shown that overexpression of macroH2A1.1 led to a more efficient iPSCs reprogramming, by not fully defined mechanisms. Cleavage under targets and tagmentation (CUT&Tag) is a recent methodology used for robust epigenomic profiling of a limited amount of cells. Here, we performed the first integrative CUT&Tag/RNA-Seq analysis of the histone variant macroH2A1-dependent orchestration of iPSCs reprogramming using human umbilical vein endothelial cells (HUVEC) during their reprogramming into iPSC over-expressing tagged macroH2A1.1 or macroH2A1.2. Our results demonstrate a higher and more widespread genome occupancy and a greater number of differentially expressed genes orchestrated by macroH2A1.1 in HUVEC undergoing reprogramming as compared to macroH2A1.2, which involved pervasive functions related to the three embryonic germ layers and increased overlap with CTCF, FOS, GATA2, and POLR2A transcription factor binding sites. In particular, all predicted macroH2A1.1 activating pathways were related to ectoderm/neural processes. As macroH2A1 isoforms have been previously associated with pathologies of the nervous system, our findings may provide relevant molecular insights for modeling neurodegenerative diseases using iPSCs.

## INTRODUCTION

Cleavage under targets and tagmentation (CUT&Tag) is a method used for efficient epigenomic profiling of small samples and single cells, devised in 2019 (Kaya-Okur et al. 2019). CUT&Tag-sequencing combines antibody-targeted controlled cleavage by a protein A-Tn5 fusion with massively parallel DNA sequencing to identify the binding sites of histone post-translational modifications (PTM), RNA Polymerase II and transcription factors (Kaya-Okur et al. 2019; Janssens et al. 2022; Bartosovic et al. 2021), at tissue scale and cellular level (Medeiros-Neto 1971). CUT&Tag can capture epigenomic heterogeneity, helping to predict sensitivity to therapeutic agents in samples from leukemic patients (Janssens et al. 2021). While Chromatin Immuno-Precipitation Sequencing (ChIP-Seq) is the most common technique utilized to study protein–DNA relations, CUT&Tag offers several advantages, such as it does not require cells to be lysed or chromatin to be fractionated, and it is suitable for single-cell platforms (Kaya-Okur et al. 2019). CUT&Tag was recently combined with RNA-seq to simultaneously profile histone PTMs and gene expression in single cells isolated from mouse brain tissue (Zhu et al. 2021). “Multiomics” approaches where the integrated analysis of transcriptional activity, histone modifications, and chromatin accessibility via CUT&Tag are still in their infancy, compared to established ChIP-seq datasets, although they have the potential to uncover the intricate epigenetic regulatory mechanisms governing different cell types.

Induced pluripotent stem cells (iPSC) are pluripotent stem cells that can be generated from a somatic cell by the introduction of four specific transcription factors (named Myc, Oct3/4, Sox2 and Klf4) key to the process (Takahashi and Yamanaka 2006). iPSCs are used in personalized disease modeling and have great potential in autologous-based regenerative medicine (Takahashi and Yamanaka 2006). However, they are generally not considered safe for transplant because of inherent iatrogenic tumorigenesis (Lee et al. 2013). There are several potential mechanisms for tumorigenicity during the induction of pluripotency in somatic cells, linked to genome and epigenome stability (Liang and Zhang 2013). The process or reprogramming involves a total remodeling of the somatic epigenetic memory, which is replaced by new iPSC-specific epigenetic patterns. Among the epigenetic alterations, the substitution of canonical histones with histone variants appears as an emerging regulator of iPSC identity (Giallongo et al. 2021b).

MacroH2A proteins are histone variants coded by 2 distinct genes: H2AFY for macroH2A1, and H2AFY2, for macroH2A2. Unlike macroH2A2, macroH2A2 is expressed ubiquitously. MacroH2A1 features two exon splicing isoforms: macroH2A1.1 and macroH2A1.2 having common and distinct biological functions in tumorigenesis, cell differentiation and stemness (Bereshchenko et al. 2019; Borghesan et al. 2016; Giallongo et al. 2020, 2021a; Lo Re et al. 2018, 2020; Lo Re and Vinciguerra 2017; Pazienza et al. 2014, 2016; Podrini et al. 2015; Rappa et al. 2013; Chiodi et al. 2021; Guberovic et al. 2022). In particular, we have recently reported that macroH2A1.1, but not macroH2A1.2, is an enhancer of DNA damage repair and iPSC reprogramming from somatic cells, increasing the efficiency of the process (Giallongo et al. 2022). Interestingly, while macroH2A1.2 does not impact iPSC differentiation into the 3 major embryonic germ layers (endoderm, mesoderm, and ectoderm), macroH2A1.1-expressing reprogrammed iPSC have a decreased potential to generate ectoderm (Giallongo et al. 2022). Understanding the genome binding and transcriptomic dynamics of macroH2A1.1 could help to improve iPSC production and availability for clinical trials. However, iPSCs reprogramming remains an inefficient process, and their amount upon somatic cell reprogramming is unsuitable for conventional ChIP-Seq approaches. In this work, we employed human umbilical vein endothelial cells (HUVEC) during their reprogramming into iPSCs over-expressing tagged macroH2A1.1 or macroH2A1.2, to perform for the first time an integrative CUT&Tag/RNA-Seq analysis of histone variant macroH2A1-dependent orchestration of human iPSC reprogramming. Our analyses uncover a new exquisite histone variant-based genomic/transcriptomic interplay underlying iPSC reprogramming, which may inform functional and preclinical assays.

## RESULTS

### Genome occupancy patterns of overexpressed macroH2A1 isoforms in HUVEC reprogrammed to iPSC

We analyzed the genome occupancy patterns of 6-His-tagged macroH2A1.1 and macroH2A1.2 in HUVEC at the 4th day of their episomal-driven reprogramming into iPSC (Giallongo et al. 2022) by using a CUT&Tag approach. Quality control performed on broad peaks suggested the exclusion of a poor-quality sequencing replicate related to macroH2A1.2 (**Supplemental_Fig_S1**). Subsequently, the IDR method was used to find consistent peaks in the remaining replicates, which, after annotation, were 12605 and 11456 for macroH2A1.1 and macroH2A1.2, respectively (**Supplemental_Table_S1 and Supplemental_Table_S2**). Their genome occupancy patterns, as well as their distances from the TSS, are plotted in **Figure 1**. The median peak sizes were 27163 and 27545 for macroH2A1.1 and macroH2A1.2 (**Supplemental_Fig_S2**). MacroH2A1.1 and macroH2A1.2 peaks were found in promoters, distal intergenic regions, and introns, with macroH2A1.2 showing a slight, not statistically significant, preference for promoters (□^2^(6,N=23953)=5.69, p=0.45) compared to macroH2A1.1. The majority of peaks in both isoforms were either near (1 Kb) or far away (> 5 Kb) from TSS.

**Figure 1.**
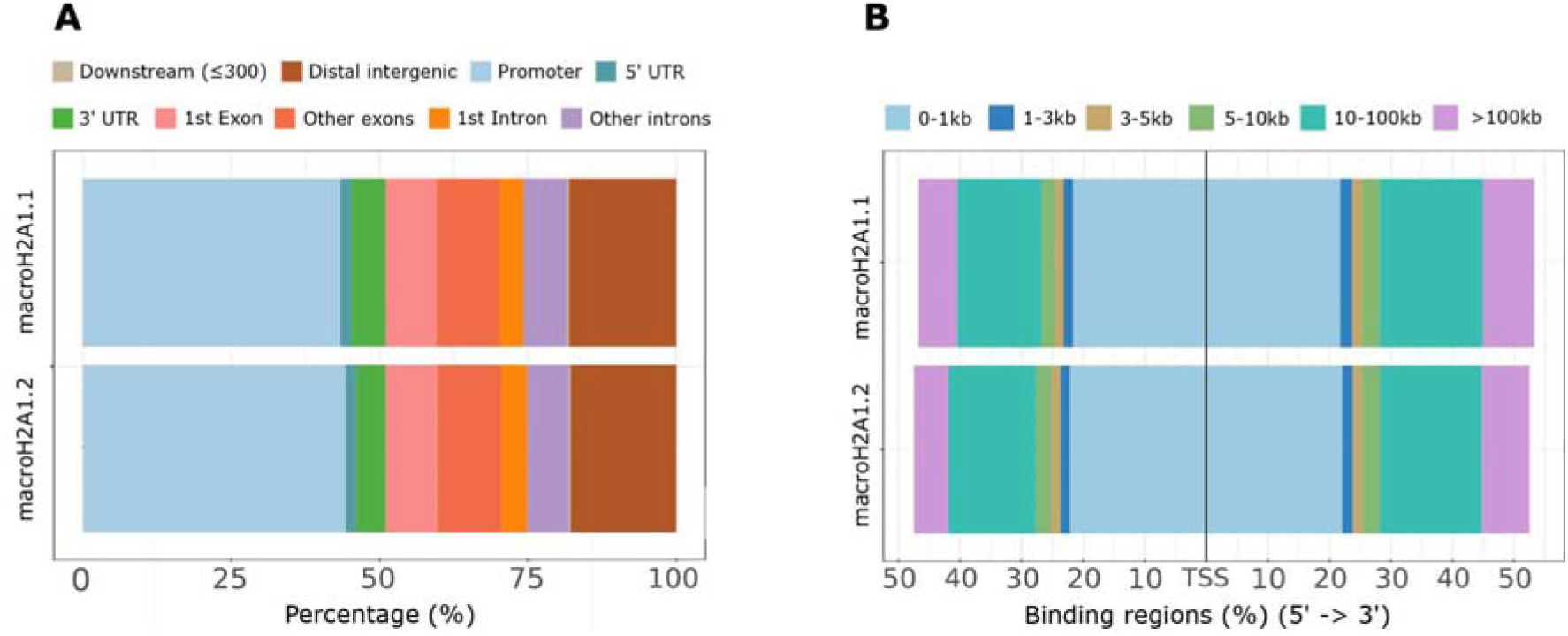
MacroH2A1.1 and macroH2A1.2 genome occupancy in relation to gene regulatory regions. Distribution of macroH2A1 isoform peaks relative to the **A)** gene structure and **B)** nearest TSS.

### Integration of CUT&Tag and transcriptomic data in HUVEC overexpressing macroH2A1 isoforms reprogrammed to iPSC

Peaks were further filtered according to whether they fell in the proximity of differentially expressed genes, as previously determined (Giallongo et al. 2022). After integration, the surviving peaks were 267 for macroH2A1.1 and 52 for macroH2A1.2. We counted 138 differentially expressed genes upon histone binding (UHB-DEG) for macroH2A1.1, of which 86 were significantly up-regulated and 52 down-regulated, and 30 UHB-DEGs for macroH2A1.2, of which 14 were up-regulated and 16 downregulated (**Supplemental_Table_S3**, **Figure 2A**). Four UHB-DEGs, i.e., *DNAH5, PPP1R37, BACH2* and *MEGF8*, were shared between the two macroH2A1 variants. The genes *DNAH5, PPP1R37, and BACH2* were overexpressed, while *MEGF8* was underexpressed by both macroH2A1.1 and macroH2A1.2 cells.

**Figure 2.**
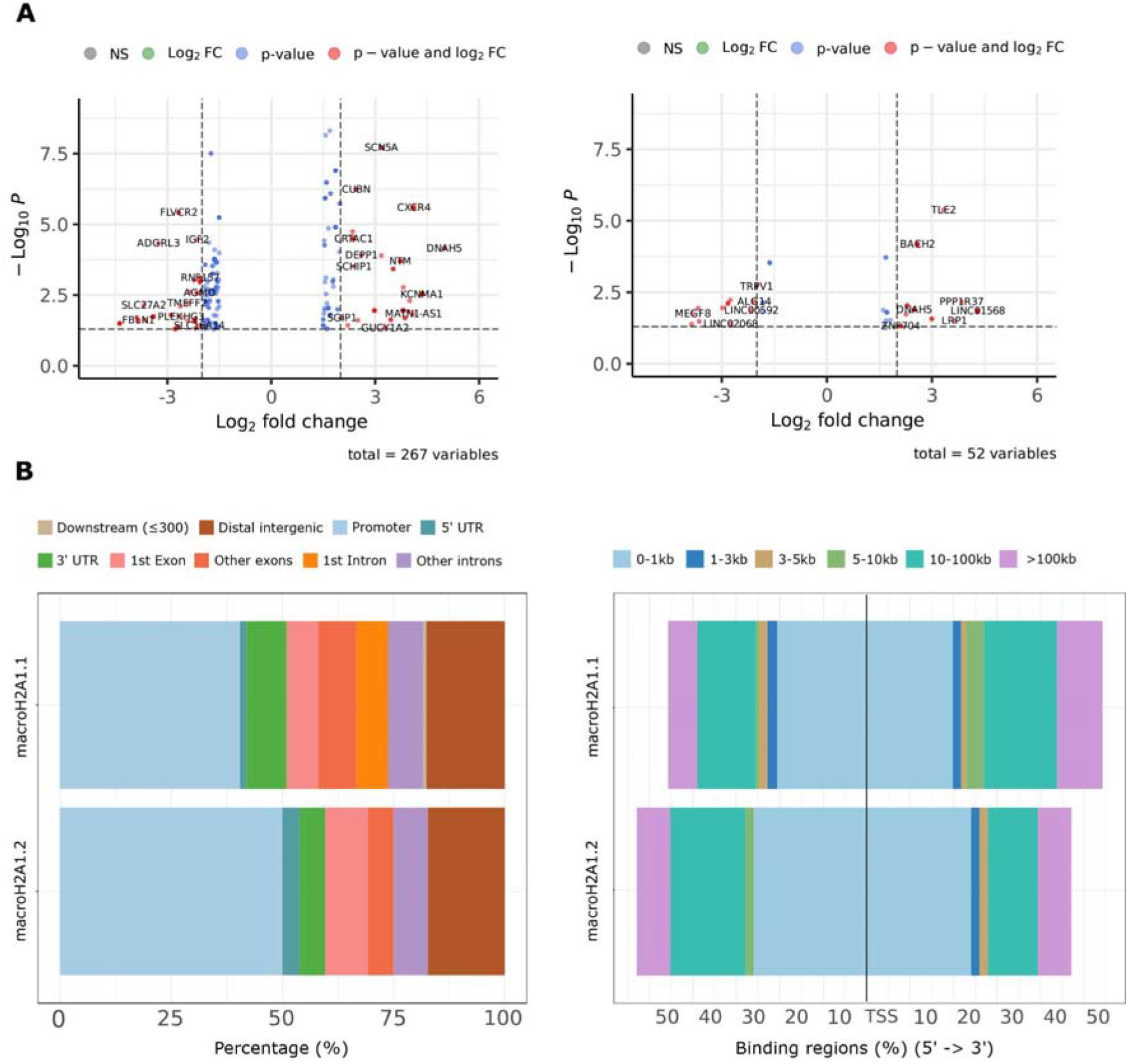
MacroH2A1.1 and macroH2A1.2 occupancy of DEGs and of their regulatory regions. **A)** Volcano plots of the significant UHB-DEGs for macroH2A1.1 (left) and macroH2A1.2 (right). **B)** Distribution of peaks in DEGs relative to the (left) gene structure and (right) distance from the nearest TSS.

The peak occupancy patterns and sizes (**Supplemental_Fig_S2**) were notably different from those in **Figure 1**. Despite the fact that macroH2A1.2’s peaks were more frequent in promoters than macroH2A1.1’s were, macroH2A1.1’s peaks were indeed more prevalent in the 3’UTR. Additionally, macroH2A1.2 did not exhibit any peaks in the first intron regions. Furthermore, compared to macroH2A1.1, macroH2A1.2’s peaks were located nearer to the TSS. The peaks of macroH2A1.2 were absent between 1 and 5 Kbp upstream and 5 to 10 Kbp downstream of the TSS **(Figure 2B)**.

### MacroH2A1 isoforms’ peaks localization within chromosomes

After integration of the genomic data obtained with CUT&tag with the RNA-Seq transcriptomic data, no peaks were found in sexual chromosomes and autosomes 18 and 21 for either isoform. MacroH2A1.1’s peaks were present in all the remaining but chromosome 16, where macroH2A1.2’s peaks were instead abundant. Additionally, macroH2A1.2’s peaks were not found on chromosomes 2, 9, 11, and 22 (**Figure 3A**). The total number of peaks identified in the chromosomes 2 (□^2^(19, N = 319) =4.67, p =3.06e-02), 5 (□^2^(19, N=319)=3.82, p =5.04e-02), 16 (□^2^(19, N=319)=25.67, p =4.04e-07), and 19 (□^2^(19, N=319)=6.65, p =9.86e-03) were significantly different between the two isoforms in HUVEC cells undergoing reprogramming. Considering the gene structure, peaks’ counts were not statistically different for any chromosome.

**Figure 3.**
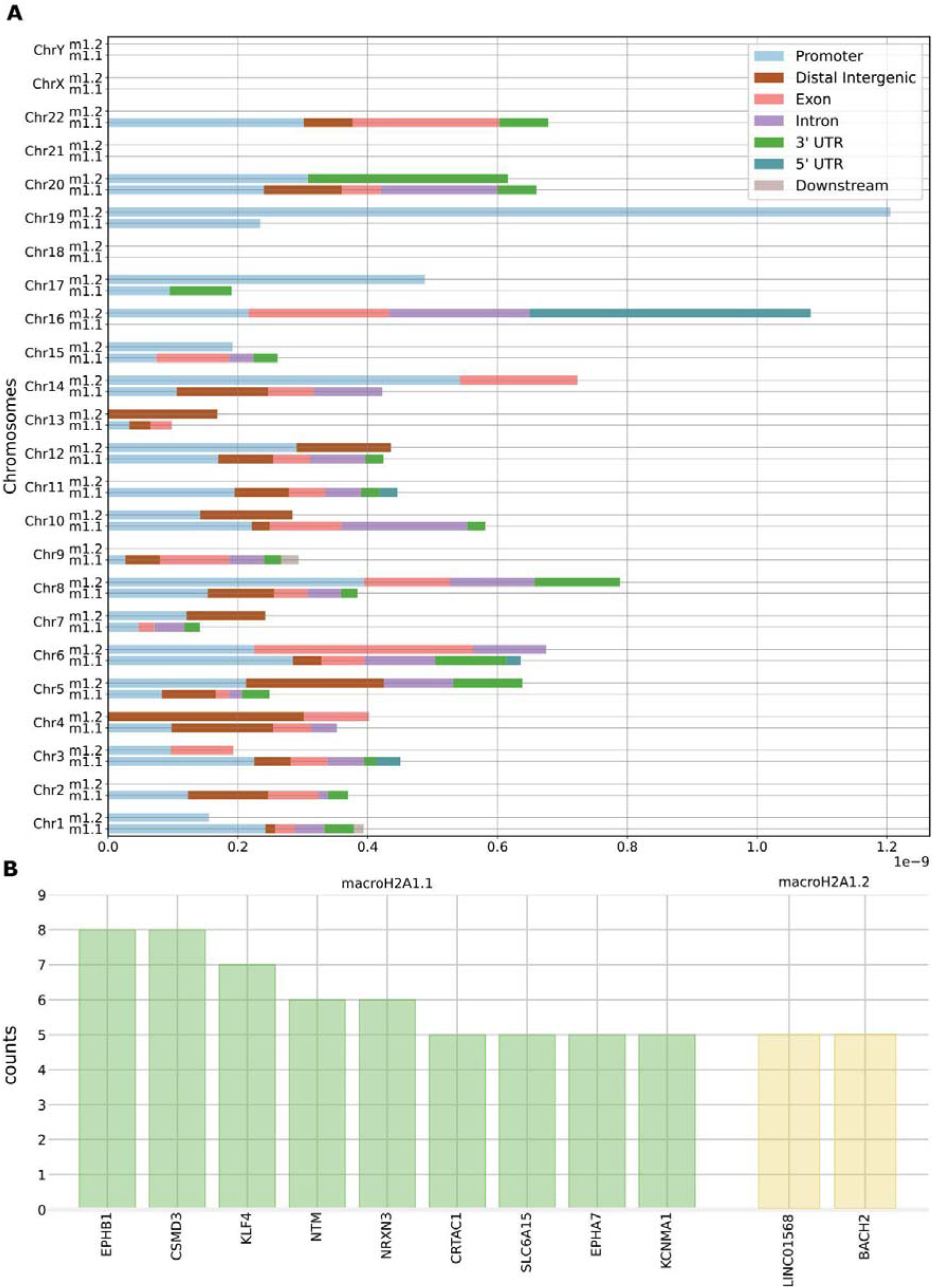
MacroH2A1.1 and macroH2A1.2 genome occupancy and regulatory regions within individual chromosomes. **A)** Normalized peak counts (X-axis) for all human chromosomes (Y-axis). **B)** Per-gene peak counts for macroH2A1.1 (left) and macroH2A1.2 (right). Threshold: per-gene peaks >4.

Moreover, multiple peaks were found in the body or proximity of a few UHB-DEGs for both isoforms. Eight peaks were found in two genes (i.e., EPHB1 and CSMD3), seven in KLF4, and six in NRXN3 and NTM for macroH2A1.1 but not for macroH2A1.2. The maximum number of peaks found in macroH2A1.2’s UHB-DEGs were, instead, five in two genes, i.e., LINC01568 and BACH2, and three peaks in five genes (DNAH5, ZNF704, MATN2, ARHGAP26, and LINC00639) (**Supplemental_Table_S4** and **S5** for macroH2A1.1 and macroH2A1.2, respectively). The UHB-DEGs shared between the macroH2A1 variants were four, i.e., *DNAH5, PPP1R37, BACH2* and *MEGF8*. Genes supported by at least 4 peaks (corresponding to the fifty percent of the highest number of peaks in a gene) are shown in **Figure 3B** (cf. also **Supplemental_Table_S4** and **S5** for macroH2A1.1 and macroH2A1.2).

### MacroH2A1.1 genome binding orchestrates expression of more genes involved in functions related to the three different germ layers than macroH2A1.2

The integrated genomic occupancy with transcriptomic data of both isoforms underwent Gene Ontology (GO) enrichment analysis. In order to emphasize the potentially distinct roles that macroH2A1.1 and macroH2A1.2 play in developmental processes, we manually annotated genes that were known to participate in the function and development of one or more of the three germ layers, i.e., ectoderm, mesoderm, and endoderm (**Supplemental_Table_S6** and **S7**). MacroH2A1.1’s peaks hit a significantly higher number of DEGs involved in biological processes related to the three different germ layers compared to macroH2A1.2, as reported in the **Supplemental_Table_S6** and **S7**. The semantic terms belonging to the Molecular Function (MF) and Cellular Component (CC) ontologies were reported in **Supplemental_Table_S8** and **S9** for macroH2A1.1 while **S10 and S11** for macroH2A1.2. Additionally, with a few exceptions, macroH2A1.2’s UHB-DEGs were not specific to a single germ layer (**Figure 4B**), while macroH2A1.1’s UHB-DEGs could be associated more specifically with one of the three germ layers (**Figure 4A**).

**Figure 4.**
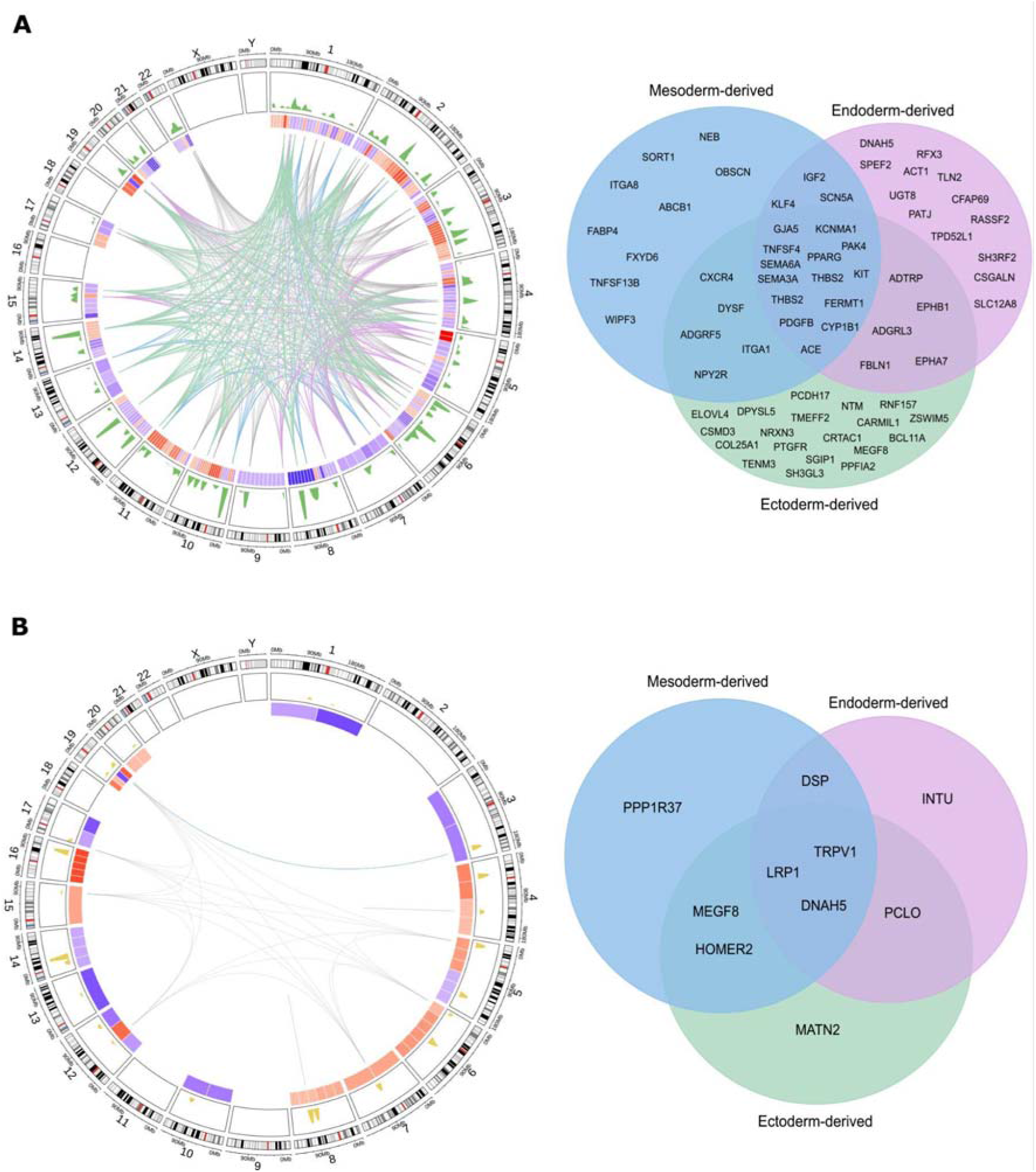
MacroH2A1 and macroH2A1.2 genome binding to DEGs involved in functions related to the three embryonic germ layers. Circos (left) and Venn diagram (right) of UHB-DEGs for **A)** macroH2A1.1 and **B)** macroH2A1.2. From the outer to the inner tracks of the circos plots: chromosomes, CUT&Tag peaks, heatmap of fold-change expression values (red and blue mean upregulated and downregulated genes in macroH2A1.1 or macroH2A1.2 overexpressing cells), genes whose biological processes relate to mesoderm (light blue links), endoderm (light purple), and ectoderm (light green). Gray links wire genes associated with more than one germ layer.

### MacroH2A1 isoform-dependent signaling pathways regulating the stemness of iPSC

The Kyoto Encyclopedia of Genes and Genomes (KEGG) was used as a starting point to study further the enrichment of macroH2A1.1 and macroH2A1.2 UHB-DEGs for biological pathways. MacroH2A1.1 was discovered to be enriched in a number of pathways, including *axon guidance* (hsa04360) and *focal adhesion* (hsa04510) (**Supplemental_Table_S12**). These pathways are known to be involved in several biological mechanisms regulating the *pluripotency of stem cells* (hsa04550), such as MAPK (hsa04010), WNT (hsa04310), PI3K (hsa04151), and *regulation of the actin cytoskeleton* (hsa04810). In contrast, macroH2A1.2 exhibited a significant enrichment in pathways that do not contribute to the regulation of stemness of iPSC (**Supplemental_Table_S13**).

UHB-DEGs were additionally enriched using Ingenuity Pathway Analysis (IPA), which is based on an orthogonal semantic knowledgebase and offers an estimate of the activation state of pathways. This was done as a cross-check to the previous analytical step. For macroH2A1.1, the total number of significant canonical pathways was 37 (**Supplemental_Fig_S4** and **Supplemental_Table_S14)**. Of these, all predicted activating pathways were related to neuronal pathways (i.e., the *semaphorin neuronal repulsive* and the *synaptogenesis* signaling pathways). Because there were so few DEGs overall, it was not possible to predict the activation state of the only two significantly enriched pathways for macroH2A1.2: *Oxytocin in Spinal Neurons* and *Inhibition of Matrix Metalloproteases* (**Supplemental_Table_S15**).

Three of the ten causal networks for different developmental processes that IPA predicted and reported in **Supplemental_Table_S16** included the development of the embryo (**Supplemental_Fig_S3A**). One of the three networks, which also described organismal and tissue development, overlapped with the majority of the other networks (**Supplemental_Fig_S3B**), demonstrating that there were numerous overlapping networks involved in the processes of embryonic development that were handled differently when macroH2A1.1 was overexpressed. However, for macroH2A1.2, only two networks could be inferred, and these networks did not represent any embryonic developmental processes (**Supplemental_Table_S16**), did not share any molecules, and were therefore disconnected.

### Regulatory signals in macroH2A1.1 and macroH2A1.2 UHB-DEGs

To gain insights into the transcriptional regulatory signals that might be mediated by macroH2A1.1 and/or macroH2A1.2 upon HUVEC reprogramming into iPCSs, we evaluated the isoforms’ capacity to mask/unmask Transcription Factor Binding Sites (TFBSs). This was accomplished by first querying for HUVEC-specific DNA regulatory features available from the ENCODE 3’s `EncRegTfbsClustered` dataset. The available features for this cell line regarded exclusively the following DNA-binding proteins: CTCF, FOS, GATA2, and POLR2A **(Supplemental_Table_S17, S18).** Among these features, 192 and 33 were located near DEGs for macroH2A1.1 and macroH2A1.2, respectively. MacroH2A1.1 and macroH2A1.2 binding sites overlapped with several CTCF responsive elements near 62 and 11 DEGs (**Supplemental_Table_S19)**. MacroH2A1.1 histone also overlapped with FOS and GATA2 binding sites near 35 and 5 DEGs, respectively; the other isoform turned out to likely influence FOS/GATA2-dependent regulation for 6 and 2 DEGs. Furthermore, the isoforms’ binding sites intersected the RNA Polymerase (POL2RA) at their promoter regions, thereby directly controlling the expression of 13 and 2 genes for macroH2A1.1 and macroH2A1.2 (**Supplemental_Table_S19)**. The expression of some genes, i.e., CUBN, KCNMA1, ZDHHC13, NRIP3, PAK, and EEF1A2, was found to be controlled by multiple bindings of macroH2A1.1 to the promoter regions of almost all CTCF, FOS, GATA2, and POL2RA DNA-binding proteins. Similarly, macroH2A1.2 appeared to regulate the expression of PPP1R37, BACH2, and ARHGAP26 genes upon binding to GATA2, FOS, and CTCF (**Supplemental_Table_S19)**.

## DISCUSSION

Previously, it was reported that macroH2A histone variants (macroH2A1 and macroH2A2) indiscriminately act as a barrier upon reprogramming towards pluripotency (Gaspar-Maia et al. 2013).

While the latter study used total knock-out cells and did not focus on macroH2A1 splicing isoforms, we reported that, in somatic cells, macroH2A1.1 overexpression activated transcriptional programs that enhanced DNA damage repair (DDR) and reprogramming. By doing so, macroH2A1.1 but not macroH2A1.2 overexpression improved iPSC reprogramming, suggesting that this splicing isoform could be a promising epigenetic target to improve iPSC genome stability and therapeutic potential (Giallongo et al. 2022).

Here, we shed light on this process: using HUVEC cells undergoing episomal reprogramming into iPSCs, we provide an integrated analysis of macroH2A1.1 and macroH2A1.2 genome occupancy and transcriptional effects, using CUT&Tag and RNA-Seq, respectively. The roles of macroH2A1 isoforms in regulating gene expression are seemingly contradictory, and the underlying mechanisms are still not well characterized. Until recently, macroH2A1 was generally believed to play a role in transcriptional repression. However, in many cases, its presence correlates also with active transcription of a subset of genes (Gamble et al. 2010; Changolkar et al. 2010; Podrini et al. 2015), as we have shown here and in our previous study (Giallongo et al. 2022). Our data clearly demonstrate that at the 4th day of reprogramming, macroH2A1.2’s DNA binding sites are more frequent in promoters and nearer to the TSS than macroH2A1.1’s are in DEGs, while macroH2A1.1’s binding sites are more prevalent in the 3’UTR and in introns. These genomic binding patterns are remarkably similar to those reported in liver cancer cells overexpressing macroH2A1.1 and macroH2A1.2 coupled to GFP, using ChIP-Seq (Hurtado-Bagès et al. 2020). These consistent differences in their genomic distribution might be reflected in the limited overlap between the genes deregulated by the different macroH2A1 isoforms in physiological and pathological processes (Borghesan et al. 2016). In breast cancer cells, macroH2A1.1 can have either an inhibitory or a stimulating effect on target gene transcription by RNA polymerase II, depending on the chromatin landscape and on differential recruitment (Recoules et al. 2022). Overall, macroH2A1.1 deposition on DEGs was more abundant and chromosome mapping revealed a more prevalent coverage compared to macroH2A1.2 in HUVEC reprogrammed to iPSCs. Interestingly, none of the macroH2A1 isoforms was present on the X chromosome of our male HUVEC (XY), which is consistent with its deposition on the inactive X chromosome in female cells, as macroH2A1 was originally identified (Bernstein et al. 2008; Mermoud et al. 1999; Chadwick and Willard 2001).

The difference between the two splice-isoforms, macroH2A1.1 and macroH2A1.2, is the usage of a mutually exclusive exon that encodes a number of amino acids defining the form and hydrophobicity of a cavity in the macrodomain (Kustatscher et al. 2005). As a consequence, macroH2A1.1, but not macroH2A1.2, is able to bind NAD+-derived ADP-ribose. Adequate activation of NAD+-dependent pathways is crucial for iPSCs pluripotency, conferring a metabolic advantage and enhancing resistance to cellular stress through kinase inhibition (Son et al. 2013; Meng et al. 2018; Teslaa and Teitell 2015). A tempting hypothesis is that enhanced sensing of NAD+-signaling mediated by macroH2A1.1 may reflect an increased efficiency of iPSCs reprogramming (Giallongo et al. 2022). During myogenic differentiation, macroH2A1.1 metabolite-binding macrodomain was essential for sustained optimal mitochondrial function but dispensable for gene regulation (Posavec Marjanoviċ et al. 2017); it remains to be elucidated whether NAD+-derived ADP-ribose binding would alter macroH2A1.1 genome occupancy in iPSCs undergoing reprogramming.

Moreover, the macroH2A1.1 binding sites hit a significantly higher number of DEGs involved in biological pathways related to the three germ layers (ectoderm, endoderm, and mesoderm) than macroH2A1.2, generating more interconnected networks; all predicted macroH2A1.1 activating pathways were related to neuronal pathways. We have previously shown that macroH2A1.1-overexpressing reprogrammed iPSCs have a decreased potential to generate the ectoderm (Giallongo et al. 2022). Interestingly, macroH2A1 levels are upregulated in tissues of patients affected by Huntington’s disease (Hu et al. 2011), and by Alzheimer’s disease (Kapadia et al. 2021). Huntington’s disease and Alzheimer’s disease human embryonic stem cells present ectodermal anomalies during development (Haremaki et al. 2019; Ryskamp et al. 2019). Moreover, systemic depletion of macroH2A1.1 in mice supports an epigenetic control necessary for increased hippocampal function and social behavior (Chiodi et al. 2021). Harnessing macroH2A1.1 in iPSC-derived neural cells may provide valuable tools for studying neurological disease mechanisms and developing potential therapies (Vadodaria et al. 2020). Our findings also show that, in HUVEC cells, macroH2A1.1 binding sites overlapped with binding sites for transcription factors CTCF, FOS, GATA2, and POLR2A to a greater extent than macroH2A1.2. These transcription factors have been consistently involved in embryonic development and in the regulation of ectoderm specification towards the nervous system in several organisms (Curran and Morgan 1995; Krendl et al. 2017; Dowen et al. 2014; Levine 2011), as well as in iPSC reprogramming (Shu et al. 2015; Song et al. 2022; Schuster et al. 2021; Markov et al. 2021). A limitation of our latter analysis is that it is restricted to HUVEC-specific DNA regulatory features present in the ENCODE 3’s `EncRegTfbsClustered` dataset. Moreover, CUT&Tag is a new technique that has not been extensively benchmarked against existing ChIP-Seq datasets on the same biological samples. A comprehensive benchmarking of CUT&Tag for H3K27ac and H3K27me3 against published ChIP-seq profiles from ENCODE in K562 cells identified a ~50% recovery of known ENCODE peaks. However, the peaks identified by CUT&Tag represent the strongest ENCODE peaks, with identical functional and biological enrichments as ChIP-seq peaks (Hu et al. 2022). We propose the multi-omics CUT&Tag and RNA-Seq complementary data integration as a powerful tool to elucidate epigenetic and gene regulatory mechanisms occurring during cell reprogramming and lineage commitment.

## METHODS

### Cell reprogramming

Human Umbilical Vein Endothelial Cells (HUVEC) were obtained from Thermo Fisher Scientific (Massachusetts, USA). Cells were cultured in Nunc EasYFlasks in Endothelial Cell Growth medium 2 (Promocell, Heidelberg, Germany). Cellular reprogramming was performed as already described (Giallongo et al. 2022). Briefly, cells were harvested using TrypLE express enzyme (Thermo Fisher Scientific, Massachusetts, USA) and transfected with Epi5 Episomal iPSC Reprogramming Kit (Thermo Fisher Scientific, Massachusetts, USA), using 10μL Neon Transfection System (Thermo Fisher Scientific, Massachusetts, USA). For this purpose, 105 living cells were resuspended in 10μL of R buffer and 1μL of the content of each vial from the reprogramming kit was added. Furthermore, 3mL of E buffer were added to the Neon transfection tube together with the vectors expressing macroH2A1.1-6His and macroH2A1.2-6His (Addgene, Massachusetts, USA), where needed [25]. Electroporation was performed using Pulse voltage 1400 V, Pulse Width 20 ms, Pulse number 2. Cells were seeded on 35mm Petri dishes coated by hESC-qualified MatriGel in endothelial medium, which was changed daily until day 4 post-transfection.

### CUT&Tag

Cells on the fourth day post-transfections were collected by TryPLE Express Enzyme (ThermoFisher, Austria) in 1.5mL tubes. CUT&Tag was performed according to the manufacturer’s protocol (Active Motif, California, USA). Briefly, cells were harvested and washed once using 1mL of 1x Wash buffer. In parallel, the Concanavalin A Beads were prepared. Each 500.000 cells, 20μL of the beads’ slurry were withdrawn and mixed with 1,6mL of 1x Binding buffer. Beads were harvested using a magnetic stand and washed twice with 1,5 mL of 1x Binding buffer. The pellet containing the cells previously harvested and the beads’ suspension were mixed in a 1.5 mL tube, rotating end-over-end for 10 minutes at room temperature. Cells were harvested by using a magnetic stand to clear and the supernatant was discarded. The pellet was resuspended in 50μL of Antibody Buffer and 1μL of Rabbit ab9108 Anti-6X His tag antibody (Abcam, Cambridge, United Kingdom) was added where needed. The mix was incubated overnight at 4oC with orbital mixing ensuring the liquid to stay together at the bottom of the tube. The Guinea Pig Anti-Rabbit Antibody secondary antibody was diluted 1:100 in Dig-Wash Buffer. The tubes were cleared using a magnetic stand and the liquid was discarded. 100μL of Guinea Pig Anti-Rabbit Antibody mix was added and tubes were incubated at room temperature for 60 minutes in an orbital rotator. Cells were cleared through a magnetic stand, and they were washed three times using 1mL of Dig-Wash Buffer. In parallel, the CUT&TAGTM Assembled pA-Tn5 Transposomes was diluted 1:100 with Dig-300 Buffer. For each immunoprecipitated sample, 100μL of CUT&TAGTM Assembled pA-Tn5 Transposomes was added, and the mix was placed on an orbital rotator for 60 minutes at room temperature. The cells were harvested using a magnetic stand and eventually washed three times using 1mL of Dig-300 buffer. In the following step each tube was supplemented with 125μL of Tagmentation Buffer and incubated at 37oC for 60 minutes. To isolate the DNA surrounding macroH2A1.1 and macroH2A1.2 deposition regions, the tagmentation was first stopped by addition of 4,2μL of 0,5M EDTA, 1,25μL of 10% SDS and 1,25μL of Proteinase K (10mg/mL) per each sample. The tubes were then vortexed at full speed and incubated for 60 minutes at 55oC to digest. The samples were cleared using a magnetic stand and 625μL of DNA Purification Buffer per each were added. The samples were transferred to DNA Purification Columns, centrifuged at 17000G for 1 minute. After the flow-through was discarded, 750μL of DNA Purification Wash Buffer was added to each sample, which was then centrifuged again at 17000G for 1 minute. The flow-through was discarded. To remove any DNA Purification Wash Buffer leftover, the empty columns were further centrifuged at 17000G for 2 minutes. To isolate the DNA, 35μL of DNA Purification Elution Buffer were added to each sample, which was then incubated at room temperature for 1 minute and eventually centrifuged at 17000G for 1 minute.

### Library Preparation and Sequencing

Library preparation for CUT&Tag samples was performed according to the manufacturer’s protocol (Active Motif, California, USA). Paired-end Illumina sequencing was obtained using Illumina Hiseq 2500 (Illumina, San Diego, CA, USA), obtaining two FASTQ files for each of the two biological replicates of macroH2A1.1, two for each of the three biological replicates of macroH2A1.2 and two for each of the three biological replicates of the control samples, for a total of 4.8, 8.1 and 7.2 million reads, respectively. The complete analytical workflow is shown in **Figure 5** and is described below.

**Figure 5.**
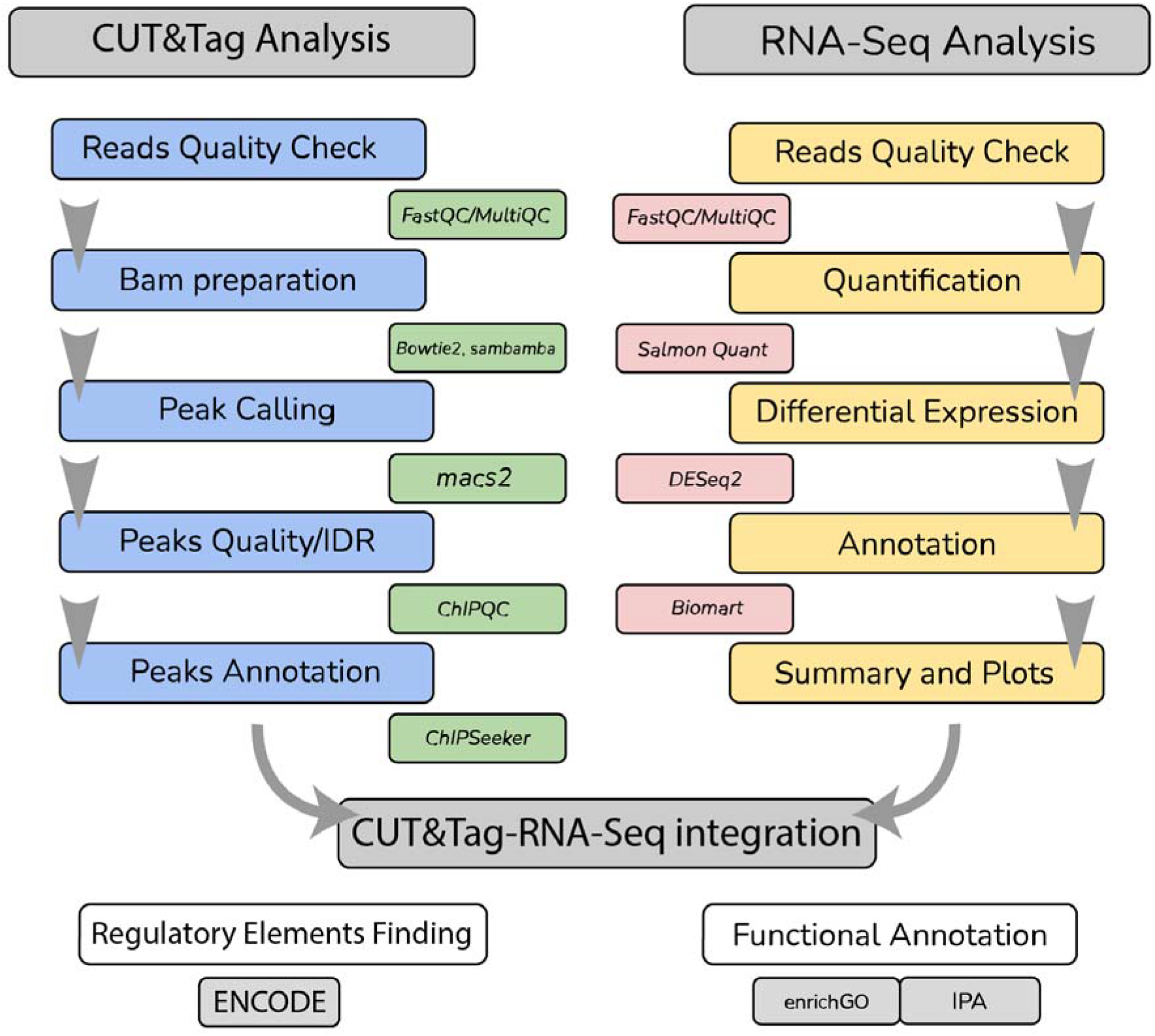
CUT&Tag and RNA-Seq integrated data analysis workflow.

### Quality-control and short-read mapping

Paired-end CUT&Tag fastq data underwent quality control analysis using the FastQC (v0.11.9) (Andrews 2010) software package and was aligned to the hg38 reference genome using Bowtie2 (v2.3.5.1) (Langmead and Salzberg 2012) with the following parameters: `-q --local --very-sensitive-local --no-mixed --no-discordant --phred33 -I 10 -X 700`. The resulting BAM files were filtered to keep only uniquely mapped reads using sambamba view (v0.8.2) (Tarasov et al. 2015), a high-performance robust tool, with the following parameters: `-F “ [XS] == null and not unmapped and not duplicate”`.

The RNA-seq data (Giallongo et al. 2022) was previously deposited into the Gene Expression Omnibus (GEO) with the dataset identifier GSE164396. Reads were quasimapped onto release 36 of the GENCODE human reference transcriptome and counted with *Salmon* v1.2.1 (Patro et al. 2017). Salmon was run with standard parameters but `validateMappings`, which was set to enable a selective alignment of the reads during mapping onto the transcriptome.

### Peak calling

Peaks were called using *MACS2* (v2.2.6) (Zhang et al. 2008) to predict the regions of the genome where histone variants were bound. Peaks were identified upon comparison between each histone variant biological replicate against a pool of baseline replicates with the following blueprint command line: `macs2 callpeak -t replicate_bam --broad -p 1e-5 -c control_bam -n sample_name -f BAMPE --keep-dup all --outdir out_dir`.

Prior to performing any downstream analyses on called peaks, a quality check was performed in R (v4.1.2) using the package *ChiPQC* v1.30.0 (Carroll et al. 2014). To measure consistency between replicates and determine an optimal cutoff for significance, we resorted to the Irreproducibility Discovery Rate (IDR) R package (v2.0.4.2) (Li et al. 2011). Peaks were annotated using the `annotatePeak` function of the *ChIPSeeker* (v1.30.3) (Yu et al. 2015) R package, referring to the TxDb.Hsapiens.UCSC.hg38.knownGene UCSC table and retrieving the Entrez Gene Identifiers of the nearest genes around the peak.

### RNA-seq data processing

Differential expression analysis was performed with *DESeq2* (Love et al. 2014), which was run with standard parameters. Genes exhibiting absolute fold-change values ≥ 1.5 and p-values <0.05 were considered differentially expressed between contrasts. The Benjamini-Hochberg procedure was used to control the multiple comparison issue.

### CUT&Tag and RNA-seq data integration

RNA-seq and CUT&Tag data were integrated using a custom Python script. The Entrez Gene Identifiers previously annotated with the *ChIPSeeker* package were converted to Ensembl Gene Identifiers using the *mygene* v3.2.2 Python module, which were then used as keys for the CUT&Tag annotated peaks and differential expression RNA-Seq data integration.

### Functional and pathway enrichment analyses

Functional annotation of CUT&Tag and RNA-seq-integrated peaks was performed using the clusterProfiler (v4.2.2) (Wu et al. 2021) R packages. The integrated data was further annotated by resorting to the following UCSC datasets: (NCBI RefSeq Functional Elements), (NCBI Transcription Factor Binding Sites), and (NCBI DNAse Hypersensitive Sites).

Functional enrichment analyses against the Gene Ontology and in particular the Biological Process (BP), Cellular Component, and Molecular Function, and the Kyoto Encyclopedia of Genes and Genomes were conducted by using the Bioconductor *enrichR* package (Xie et al. 2021). Functions and pathways were considered significantly over-represented if their p-values were <0.05.

As a crosscheck and in order to estimate the activation state of enriched pathways, we used Ingenuity Pathway Analysis (IPA). IPA is based on the calculation of the *activation z score* as a way to infer the activation states of predicted transcriptional regulators. Positive and negative z-scores are associated with activated and inhibited pathways, respectively. The more extreme a z-score, the more statistically confident a prediction is (Krämer et al. 2014). IPA was finally used to determine the ten most likely causal networks, where upstream molecules were also considered in order to expose causal relationships associated with our experimental data and confirm the most represented pathways.

All results were plotted using the *ggplot2* (v.3.3.5), *enrichPlot* (v.1.14.2), and *circlize* (v.0.4.15) R packages.

### Histones’ binding sites annotation

Binding sites near DEGs were evaluated for the presence of known regulatory elements. The genomic coordinates of all the elements reported in the `encRegTfbsClustered` dataset, a compilation of Transcription Factor ChIP-seq Clusters relative to 340 factors and 129 cell types from ENCODE 3 (ENCODE 3 - encRegTfbsClustered), were intersected with the peaks’ chromosomal coordinates by considering a minimum of 1 bp overlap. Overlapping regulatory sites with a score ≥900, which is calculated at UCSC ranging from 1 to 1000 and is based on signal values assigned by the ENCODE pipeline, and experimental source `=endothelial_cell_of_umbilical_vein` were retained.

### Statistical analysis

The module `chi2_contingency` of the *scipy.stats* (v1.7.3) Python package was used to test the differential distribution of peak frequency between chromosomes. One-way chisquare test was performed using the `chisquare` module of *scipy.stats*. The tests were considered significant whenever the p-value <0.05. The barplot of chromosomes was drawn by considering the normalized peak frequencies:

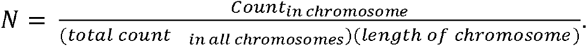

## DATA ACCESS

The RNA-seq data (Giallongo et al. 2022) was previously deposited into the Gene Expression Omnibus (GEO) with the dataset identifier GSE164396. Genomics sequences obtained in this study by CUT&Tag have been deposited into the GEO with the dataset identifier GSE214013.

## ACKNOWLEDGMENTS

This research was funded by the European Regional Development Fund-Project MAGNET (CZ.02.1.01/0.0/0.0/15_003/0000492), the Ministry of Health of the Czech Republic (NV18-03-00058), the Italian Ministry of Health and ‘5×1000’ voluntary contributions (R22-5×1000), and by the European Commission Horizon 2020 Framework Program (project 856871— TRANSTEM). We thank Genomix4Life (Baronissi, Italy) and CEITEC (Brno, Czech Republic) for excellent technical assistance with genomics and transcriptomics approaches, respectively.

## REFERENCES

Andrews S. 2010. FastQC: A Quality Control Tool for High Throughput Sequence Data. FastQC. http://www.bioinformatics.babraham.ac.uk/projects/fastqc/ (Accessed September 19, 2022).

Bartosovic M, Kabbe M, Castelo-Branco G. 2021. Single-cell CUT&Tag profiles histone modifications and transcription factors in complex tissues. Nat Biotechnol 39: 825–835.

Bereshchenko O, Lo Re O, Nikulenkov F, Flamini S, Kotaskova J, Mazza T, Le Pannérer MM, Buschbeck M, Giallongo C, Palumbo G, et al. 2019. Deficiency and haploinsufficiency of histone macroH2A1.1 in mice recapitulate hematopoietic defects of human myelodysplastic syndrome. Clin Epigenetics 11: 121.

Bernstein E, Muratore-Schroeder TL, Diaz RL, Chow JC, Changolkar LN, Shabanowitz J, Heard E, Pehrson JR, Hunt DF, Allis CD. 2008. A phosphorylated subpopulation of the histone variant macroH2A1 is excluded from the inactive X chromosome and enriched during mitosis. Proc Natl Acad Sci U S A 105: 1533–1538.

Borghesan M, Fusilli C, Rappa F, Panebianco C, Rizzo G, Oben JA, Mazzoccoli G, Faulkes C, Pata I, Agodi A, et al. 2016. DNA Hypomethylation and Histone Variant macroH2A1 Synergistically Attenuate Chemotherapy-Induced Senescence to Promote Hepatocellular Carcinoma Progression. Cancer Res 76: 594–606.

Carroll TS, Liang Z, Salama R, Stark R, de Santiago I. 2014. Impact of artifact removal on ChIP quality metrics in ChIP-seq and ChIP-exo data. Front Genet 5: 75.

Chadwick BP, Willard HF. 2001. Histone H2A variants and the inactive X chromosome: identification of a second macroH2A variant. Hum Mol Genet 10: 1101–1113.

Changolkar LN, Singh G, Cui K, Berletch JB, Zhao K, Disteche CM, Pehrson JR. 2010. Genome-wide distribution of macroH2A1 histone variants in mouse liver chromatin. Mol Cell Biol 30: 5473–5483.

Chiodi V, Domenici MR, Biagini T, De Simone R, Tartaglione AM, Di Rosa M, Lo Re O, Mazza T, Micale V, Vinciguerra M. 2021. Systemic depletion of histone macroH2A1.1 boosts hippocampal synaptic plasticity and social behavior in mice. FASEB J 35: e21793.

Curran T, Morgan JI. 1995. Fos: an immediate-early transcription factor in neurons. J Neurobiol 26: 403–412.

Dowen JM, Fan ZP, Hnisz D, Ren G, Abraham BJ, Zhang LN, Weintraub AS, Schujiers J, Lee TI, Zhao K, et al. 2014. Control of cell identity genes occurs in insulated neighborhoods in mammalian chromosomes. Cell 159: 374–387.

ENCODE 3 - encRegTfbsClustered. encRegTfbsClustered. http://hgdownload.cse.ucsc.edu/goldenpath/hg19/encRegTfbsClustered/ (Accessed September 15, 2022).

Gamble MJ, Frizzell KM, Yang C, Krishnakumar R, Kraus WL. 2010. The histone variant macroH2A1 marks repressed autosomal chromatin, but protects a subset of its target genes from silencing. Genes Dev 24: 21–32.

Gaspar-Maia A, Qadeer ZA, Hasson D, Ratnakumar K, Leu NA, Leroy G, Liu S, Costanzi C, Valle-Garcia D, Schaniel C, et al. 2013. MacroH2A histone variants act as a barrier upon reprogramming towards pluripotency. Nat Commun 4: 1565.

Giallongo S, Di Rosa M, Caltabiano R, Longhitano L, Reibaldi M, Distefano A, Lo Re O, Amorini AM, Puzzo L, Salvatorelli L, et al. 2020. Loss of macroH2A1 decreases mitochondrial metabolism and reduces the aggressiveness of uveal melanoma cells. Aging 12: 9745–9760.

Giallongo S, Lo Re O, Lochmanová G, Parca L, Petrizzelli F, Zdráhal Z, Mazza T, Vinciguerra M. 2021a. Phosphorylation within Intrinsic Disordered Region Discriminates Histone Variant macroH2A1 Splicing Isoforms-macroH2A1.1 and macroH2A1.2. Biology 10. http://dx.doi.org/10.3390/biology10070659.

Giallongo S, Řeháková D, Biagini T, Lo Re O, Raina P, Lochmanová G, Zdráhal Z, Resnick I, Pata P, Pata I, et al. 2022. Histone Variant macroH2A1.1 Enhances Nonhomologous End Joining-dependent DNA Double-strand-break Repair and Reprogramming Efficiency of Human iPSCs. Stem Cells 40: 35–48. http://dx.doi.org/10.1093/stmcls/sxab004.

Giallongo S, Rehakova D, Raffaele M, Lo Re O, Koutna I, Vinciguerra M. 2021b. Redox and Epigenetics in Human Pluripotent Stem Cells Differentiation. Antioxid Redox Signal 34: 335–349.

Guberovic I, Farkas M, Corujo D, Buschbeck M. 2022. Evolution, structure and function of divergent macroH2A1 splice isoforms. Semin Cell Dev Biol. http://dx.doi.org/10.1016/j.semcdb.2022.03.036.

Haremaki T, Metzger JJ, Rito T, Ozair MZ, Etoc F, Brivanlou AH. 2019. Self-organizing neuruloids model developmental aspects of Huntington’s disease in the ectodermal compartment. Nat Biotechnol 37: 1198–1208.

Hu D, Abbasova L, Schilder BM, Nott A, Skene NG, Marzi SJ. 2022. CUT&Tag recovers up to half of ENCODE ChIP-seq peaks. bioRxiv. http://biorxiv.org/lookup/doi/10.1101/2022.03.30.486382.

Hurtado-Bagès S, Posavec Marjanovic M, Valero V, Malinverni R, Corujo D, Bouvet P, Lavigne A-C, Bystricky K, Buschbeck M. 2020. The Histone Variant MacroH2A1 Regulates Key Genes for Myogenic Cell Fusion in a Splice-Isoform Dependent Manner. Cells 9. http://dx.doi.org/10.3390/cells9051109.

Hu Y, Chopra V, Chopra R, Locascio JJ, Liao Z, Ding H, Zheng B, Matson WR, Ferrante RJ, Rosas HD, et al. 2011. Transcriptional modulator H2A histone family, member Y (H2AFY) marks Huntington disease activity in man and mouse. Proc Natl Acad Sci U S A 108: 17141–17146.

Janssens DH, Meers MP, Wu SJ, Babaeva E, Meshinchi S, Sarthy JF, Ahmad K, Henikoff S. 2021. Automated CUT&Tag profiling of chromatin heterogeneity in mixed-lineage leukemia. Nat Genet 53: 1586–1596.

Janssens DH, Otto DJ, Meers MP, Setty M, Ahmad K, Henikoff S. 2022. CUT&Tag2for1: a modified method for simultaneous profiling of the accessible and silenced regulome in single cells. Genome Biol 23: 81.

Kapadia M, Mian MF, Ma D, Hutton CP, Azam A, Narkaj K, Cao C, Brown B, Michalski B, Morgan D, et al. 2021. Differential effects of chronic immunosuppression on behavioral, epigenetic, and Alzheimer’s disease-associated markers in 3xTg-AD mice. Alzheimers Res Ther 13: 30.

Kaya-Okur HS, Wu SJ, Codomo CA, Pledger ES, Bryson TD, Henikoff JG, Ahmad K, Henikoff S. 2019. CUT&Tag for efficient epigenomic profiling of small samples and single cells. Nat Commun 10: 1930.

Krämer A, Green J, Pollard J Jr, Tugendreich S. 2014. Causal analysis approaches in Ingenuity Pathway Analysis. Bioinformatics 30: 523–530.

Krendl C, Shaposhnikov D, Rishko V, Ori C, Ziegenhain C, Sass S, Simon L, Müller NS, Straub T, Brooks KE, et al. 2017. GATA2/3-TFAP2A/C transcription factor network couples human pluripotent stem cell differentiation to trophectoderm with repression of pluripotency. Proc Natl Acad Sci U S A 114: E9579–E9588.

Kustatscher G, Hothorn M, Pugieux C, Scheffzek K, Ladurner AG. 2005. Splicing regulates NAD metabolite binding to histone macroH2A. Nat Struct Mol Biol 12: 624–625.

Langmead B, Salzberg SL. 2012. Fast gapped-read alignment with Bowtie 2. Nat Methods 9: 357–359.

Lee AS, Tang C, Rao MS, Weissman IL, Wu JC. 2013. Tumorigenicity as a clinical hurdle for pluripotent stem cell therapies. Nat Med 19: 998–1004.

Levine M. 2011. Paused RNA polymerase II as a developmental checkpoint. Cell 145: 502–511.

Liang G, Zhang Y. 2013. Genetic and epigenetic variations in iPSCs: potential causes and implications for application. Cell Stem Cell 13: 149–159.

Li Q, Brown JB, Huang H, Bickel PJ. 2011. Measuring reproducibility of high-throughput experiments. Ann Appl Stat 5: 1752–1779.

Lo Re O, Douet J, Buschbeck M, Fusilli C, Pazienza V, Panebianco C, Castracani CC, Mazza T, Li Volti G, Vinciguerra M. 2018. Histone variant macroH2A1 rewires carbohydrate and lipid metabolism of hepatocellular carcinoma cells towards cancer stem cells. Epigenetics 13: 829–845.

Lo Re O, Mazza T, Giallongo S, Sanna P, Rappa F, Vinh Luong T, Li Volti G, Drovakova A, Roskams T, Van Haele M, et al. 2020. Loss of histone macroH2A1 in hepatocellular carcinoma cells promotes paracrine-mediated chemoresistance and CD4CD25FoxP3 regulatory T cells activation. Theranostics 10: 910–924.

Lo Re O, Vinciguerra M. 2017. Histone MacroH2A1: A Chromatin Point of Intersection between Fasting, Senescence and Cellular Regeneration. Genes 8. http://dx.doi.org/10.3390/genes8120367.

Love MI, Huber W, Anders S. 2014. Moderated estimation of fold change and dispersion for RNA-seq data with DESeq2. Genome Biol 15: 550.

Markov GJ, Mai T, Nair S, Shcherbina A, Wang YX, Burns DM, Kundaje A, Blau HM. 2021. AP-1 is a temporally regulated dual gatekeeper of reprogramming to pluripotency. Proc Natl Acad Sci U S A 118. http://dx.doi.org/10.1073/pnas.2104841118.

Medeiros-Neto GA. 1971. Respiration and iodine transport by thyroid slices as influenced by ouabain, succinate, -ketoglutarate, sodium and potassium. Acta Physiol Lat Am 21: 126–136.

Meng Y, Ren Z, Xu F, Zhou X, Song C, Wang VY-F, Liu W, Lu L, Thomson JA, Chen G. 2018. Nicotinamide Promotes Cell Survival and Differentiation as Kinase Inhibitor in Human Pluripotent Stem Cells. Stem Cell Reports 11: 1347–1356.

Mermoud JE, Costanzi C, Pehrson JR, Brockdorff N. 1999. Histone macroH2A1.2 relocates to the inactive X chromosome after initiation and propagation of X-inactivation. J Cell Biol 147: 1399–1408.

NCBI DNAse Hypersensitive Sites. DNAse hypersensitive Sites. https://hgdownload.soe.ucsc.edu/gbdb/hg38/bbi/wgEncodeRegDnase/wgEncodeRegDnaseUwHuvecHotspot.broadPeak.bb (Accessed September 15, 2022).

NCBI RefSeq Functional Elements. NCBI RefSeq Functional Elements. http://hgdownload.soe.ucsc.edu/gbdb/hg38/ncbiRefSeq/refSeqFuncElems.bb (Accessed September 15, 2022).

NCBI Transcription Factor Binding Sites. Transcription Factor Binding Sites. http://hgdownload.soe.ucsc.edu/goldenPath/hg38/encRegTfbsClustered/encRegTfbsClusteredWithCells.hg38.bed.gz (Accessed September 15, 2022).

Patro R, Duggal G, Love MI, Irizarry RA, Kingsford C. 2017. Salmon provides fast and bias-aware quantification of transcript expression. Nat Methods 14: 417–419.

Pazienza V, Borghesan M, Mazza T, Sheedfar F, Panebianco C, Williams R, Mazzoccoli G, Andriulli A, Nakanishi T, Vinciguerra M. 2014. SIRT1-metabolite binding histone macroH2A1.1 protects hepatocytes against lipid accumulation. Aging 6: 35–47.

Pazienza V, Panebianco C, Rappa F, Memoli D, Borghesan M, Cannito S, Oji A, Mazza G, Tamburrino D, Fusai G, et al. 2016. Histone macroH2A1.2 promotes metabolic health and leanness by inhibiting adipogenesis. Epigenetics Chromatin 9: 45.

Podrini C, Koffas A, Chokshi S, Vinciguerra M, Lelliott CJ, White JK, Adissu HA, Williams R, Greco A. 2015. MacroH2A1 isoforms are associated with epigenetic markers for activation of lipogenic genes in fat-induced steatosis. FASEB J 29: 1676–1687.

Posavec Marjanoviċ M, Hurtado-Bagès S, Lassi M, Valero V, Malinverni R, Delage H, Navarro M, Corujo D, Guberovic I, Douet J, et al. 2017. MacroH2A1.1 regulates mitochondrial respiration by limiting nuclear NAD consumption. Nat Struct Mol Biol 24: 902–910.

Rappa F, Greco A, Podrini C, Cappello F, Foti M, Bourgoin L, Peyrou M, Marino A, Scibetta N, Williams R, et al. 2013. Immunopositivity for histone macroH2A1 isoforms marks steatosis-associated hepatocellular carcinoma. PLoS One 8: e54458.

Recoules L, Heurteau A, Raynal F, Karasu N, Moutahir F, Bejjani F, Jariel-Encontre I, Cuvier O, Sexton T, Lavigne A-C, et al. 2022. The histone variant macroH2A1.1 regulates RNA polymerase II-paused genes within defined chromatin interaction landscapes. J Cell Sci 135. http://dx.doi.org/10.1242/jcs.259456.

Ryskamp DA, Korban S, Zhemkov V, Kraskovskaya N, Bezprozvanny I. 2019. Neuronal Sigma-1 Receptors: Signaling Functions and Protective Roles in Neurodegenerative Diseases. Front Neurosci 13: 862.

Schuster J, de Guidi C, Fatima A, Sobol M, Dahl N. 2021. Syndromic RNA polymerase II insufficiency: Generation of a human induced pluripotent stem cell line (UUIGPi002A-5) with a heterozygous disruption of POLR2A. Stem Cell Res 57: 102577.

Shu J, Zhang K, Zhang M, Yao A, Shao S, Du F, Yang C, Chen W, Wu C, Yang W, et al. 2015. GATA family members as inducers for cellular reprogramming to pluripotency. Cell Res 25: 169–180.

Song Y, Liang Z, Zhang J, Hu G, Wang J, Li Y, Guo R, Dong X, Babarinde IA, Ping W, et al. 2022. CTCF functions as an insulator for somatic genes and a chromatin remodeler for pluripotency genes during reprogramming. Cell Rep 39: 110626.

Son MJ, Son M-Y, Seol B, Kim M-J, Yoo CH, Han M-K, Cho YS. 2013. Nicotinamide overcomes pluripotency deficits and reprogramming barriers. Stem Cells 31: 1121–1135.

Takahashi K, Yamanaka S. 2006. Induction of pluripotent stem cells from mouse embryonic and adult fibroblast cultures by defined factors. Cell 126: 663–676.

Tarasov A, Vilella AJ, Cuppen E, Nijman IJ, Prins P. 2015. Sambamba: fast processing of NGS alignment formats. Bioinformatics 31: 2032–2034.

Teslaa T, Teitell MA. 2015. Pluripotent stem cell energy metabolism: an update. EMBO J 34: 138–153.

Vadodaria KC, Jones JR, Linker S, Gage FH. 2020. Modeling Brain Disorders Using Induced Pluripotent Stem Cells. Cold Spring Harb Perspect Biol 12. http://dx.doi.org/10.1101/cshperspect.a035659.

Wu T, Hu E, Xu S, Chen M, Guo P, Dai Z, Feng T, Zhou L, Tang W, Zhan L, et al. 2021. clusterProfiler 4.0: A universal enrichment tool for interpreting omics data. Innovation (Camb) 2: 100141.

Xie Z, Bailey A, Kuleshov MV, Clarke DJB, Evangelista JE, Jenkins SL, Lachmann A, Wojciechowicz ML, Kropiwnicki E, Jagodnik KM, et al. 2021. Gene Set Knowledge Discovery with Enrichr. Curr Protoc 1: e90.

Yu G, Wang L-G, He Q-Y. 2015. ChIPseeker: an R/Bioconductor package for ChIP peak annotation, comparison and visualization. Bioinformatics 31: 2382–2383.

Zhang Y, Liu T, Meyer CA, Eeckhoute J, Johnson DS, Bernstein BE, Nusbaum C, Myers RM, Brown M, Li W, et al. 2008. Model-based analysis of ChIP-Seq (MACS). Genome Biol 9: R137.

Zhu C, Zhang Y, Li YE, Lucero J, Behrens MM, Ren B. 2021. Joint profiling of histone modifications and transcriptome in single cells from mouse brain. Nat Methods 18: 283–292.

